# A Na_2_CO_3_-responsive chitinase gene from *Chinese wildrye* improve pathogen resistance and saline-alkali stress tolerance in transgenic tobacco and maize

**DOI:** 10.1101/707281

**Authors:** Xiang-Guo Liu, Ying Yu, Qing Liu, Suren Deng, Xue-Bo Jin, Yue-Jia Yin, Jia Guo, Nan Li, Yang Liu, Si-ping Han, Chuang Wang, Dong-Yun Hao

## Abstract

Salinity and microbial pathogens are the major limiting factors for crop production. Although the manipulation of many genes could improve plant performance under either of these stresses, few genes have reported that could improve both pathogen resistance and saline-alkali stress tolerance. In this study, we identified a new chitinase gene *<u>CHI</u>TINASE 2* (*LcCHI2*) that encodes a class II chitinase from a Chinese wildrye (*Leymus Chinensis*), which grows naturally on alkaline-sodic soil. Overexpression of *LcCHI2* increased chitinase activity in transgenic plants. The transgenic tobacco and maize exhibited improved pathogen resistance and enhanced both neutral salt and alkaline salt stress tolerance. Overexpression of *LcCHI2* reduced sodium (Na^+^) accumulation, malondialdehyde content and relative electrical conductivity in transgenic tobacco under salt stress. In addition, the transgenic tobacco showed diminished lesion against bacterial and fungal pathogen challenge, suggesting an improved disease resistance. Similar improved performance was also observed in *LcCHI2*-overexpressed maize under both pathogen and salt stresses. It is worth noting that this genetic manipulation does not impair the growth and yield of transgenic tobacco and maize under normal cultivation condition. Apparently, application of *LcCHI2* provides a new train of thought for genetically engineering saline-alkali and pathogen resistant crops of both dicots and monocots.

## Introduction

The biotic and abiotic stresses, such as pathogens and salinity, severely affected crop growth and agricultural productivity worldwide. According to incomplete statistics of UNESCO and FAO, 950 million ha (6.4%) of the world’s land area has saline-alkali soil, and about 10% of this area is found in China. The alkaline soils of China contains high levels of Na_2_CO_3_ and NaHCO_3_ (Wang *et al.*, 1993). Only few species of plants could grow in such soils and they are sparsely distributed. Chinese wildrye (*Leymus Chinensis*), a perennial grass of family gramineae, grows naturally on alkaline soil, suggesting the existence of such a mechanism. Apparently, identification of functional genes specifically responsible for stress tolerance would be a prerequisite of crop improvement through biotech breeding.

Chitinase is one of the important enzyme family that has been found to involve in such a mechanism of divergent biological functions. Chitinases (EC 3.2.1.14) are a family of glycosyl hydrolases (GHs) responsible for the hydrolysis of the chitin polymer (a β-1,4-linked *N*-acetylglucosamine), a structural component found in the cell walls of fungi, insects, a variety of crustaceans and nematode eggs (Kesari *et al.*, 2015). Interestingly, although the chitin is not present in plants, different subgroups of chitinase genes were identified in plants (Kesari *et al.*, 2015). Plant chitinase proteins are generally divided into six classes, I to VI (Patil *et al.*, 2000). The classes III and V belong to the GH18 family, whereas class I, II, IV and VI belong to the GH19 family (Patil *et al.*, 2000). Both families exhibit diversity in their nucleic acid sequences, protein structures, substrate specificities, sensitivity to inhibitors and mechanisms of catalysis. Along with active chitinases, the plant genome also consists of a large number of sequences encoding catalytically inactive chitinases that are referred to as chitinase-like (CTL) proteins (Kesari *et al.*, 2015). A few studies suggested that the likely substrates of plant CTL proteins may be arabinogalactan protein, chitooligosaccharides, *N*-acetylchitooligosaccharides and other GlcNAc-containing glycoproteins or glycolipids, but the substrates of most plant CTL proteins remain uncertain (Sanchez-Rodriguez *et al.*, 2012; Zhong *et al.*, 2002; van Hengel *et al.*, 2001). The intrinsic diversities of plant chitinase imply that the chitinase/CTL genes likely have a broad function.

Because chitin is the major component of fungal cell walls, earlier studies on the role of chitinase genes focused extensively on its involvement in plant defense responses to fungal pathogen infection (Verburg and Huynh, 1991; Brogue *et al.*, 1991). Chitinase gene expression in plant tissues is strongly induced by fungal pathogens and chitin oligosaccharides (Seo *et al.*, 2008; Hong and Hwang, 2002). Interestingly, the expression of chitinase gene responded also to infection of viruses, bacteria and oomycetes, which do not have chitin or related structures in their cell wall (Hong and Hwang, 2006; van Loon *et al.*, 2006). Accordingly, a number of plant chitinases are defined as classic pathogenesis-related proteins (PRs) including PR-3, -4, -8 and -11 families (van Loon *et al.*, 2006). Overexpression of these PR proteins conferred resistance to plant disease in different plant species (Brogue *et al.*, 1991; Yamamoto *et al.*, 2000; Tabaeizadeh *et al.*, 1999; Nishizawa *et al.*, 1999; Shin *et al.*, 2008; Takahashi *et al.*, 2005).

Plants chitinase and CTL genes play a role not only in defense related processes but also in abiotic stress tolerance. In *Arabidopsis thaliana*, the *AtPR3* gene which encodes a class II chitinase was induced by high salt (Seo *et al.*, 2008). The pepper class II basic chitinase gene *CaChi2* was induced by salt, drought and osmotic stresses (Hong and Hwang, 2002, 2006). The expression of *ptch28* from *Lycopersicon chilense* plants induced by osmotic and abscisic acid (Chen *et al.*, 1994). In addition to salt, osmotic and drought stresses, expression of the class II chitinase gene from Bermuda grass was reported to be up-regulated by cold stress and function as an antifreeze protein (Nakamura *et al.*, 2008). Similarly, in winter rye (*Secale cereal*) and highbush blueberry, cold stress induced the chitinase like proteins, which showed antifreeze activity *in vitro* (Yeh *et al.*, 2000; Kikuchi and Masuda, 2009). Interestingly, most of the reported chitinase genes that responsive to both biotic and abiotic stresses belonged to the class II family and homologue to PR3 proteins (Figure S1). Furthermore, overexpression of these PR3 genes, such as *AtPR3* and *CaChi2*, significantly increased the salt resistance of the transgenic plants (Seo *et al.*, 2008; Hong and Hwang, 2006). In contrast, mutation of the *AtCTL1* resulted in oversensitive to salt and drought stresses (Kwon *et al.*, 2007).

Wildly grown plants have developed through evolution into adaptive mechanisms against the various environmental threatening. Apparently, identification of functional genes specifically responsible for stress tolerance would be a prerequisite of crop improvement through biotech breeding. In a previous report, a chitinase like EST sequence was isolated from Chinese wildrye which was treated with Na_2_CO_3_ (Jin *et al.*, 2006). Accordingly, we cloned a unique chitinase gene and designated it as *LcCHI2*, which belongs to class II chitinase gene family. In the cultivated wildrye, expression of *LcCHI2* was induced by salt stress leading to an increase in chitinase activity. Overexpression of *LcCHI2* increased pathogen and salt stress resistance in both tobacco and Maize. This study provides a novel train of thought in improvement of both pathogen resistance and salt stress tolerance, in particular alkaline salt stress, through overexpression of a single gene in dicots and monocots.

## Material and methods

### Cloning and expression analysis of *LcCHI2*

Total RNA was extracted from 4-week-old seedlings of Chinese wildrye treated with 100 mM Na_2_CO_3_ using RNAiso plus (TaKaRa, Japan) according to the instruction manual. Full-length cDNA of LcCHI2 sequence was obtained using a SMART™ RACE cDNA Amplification Kit (Clontech, USA) for 3’ RACE with the primers 5′-CCGACCAGTTCCAATGGGGCT-3′ and 5′-GGCCACGCGTCGACTAGTACTTTTTTTTTTTTTTTTT-3′.

For Expression analysis of *LcCHI2*, total RNA was extracted from 4-week-old Chinese wildrye harvested at different time points under various stress treatments. A 771 bp fragment of *LcCHI2* cDNA was amplified in 30 cycles with the primers 5′-ATGGCGAGGTTTGCTGCCCTCG-3′ and 5′-CTAGCTAGCGAAGTTTCGCTGGGTG-3′. As an internal standard for cDNA levels, a 245 bp fragment of ACTIN cDNA (GenBank accession number AB181991) was amplified simultaneously in 28 cycles with primers 5′-GGACCTTGCTGGTCGTGACC-3′and 5′-CCTCAGGGCACCTGAACCTTT-3′. DIG High Prime DNA Labeling and Detection Starter Kit I (Roche, USA) was used for Northern blot analysis. The specific *LcCHI2* cDNA fragment was amplified with the primers 5′-ATGGCGAGGTTTGCTGCCC-3′ and 5′-CTAGCTAGCGAAGTTTCGCTGGG-3′ and then labeled with DIG-High Prime used for the hybridization probe.

### Agrobacterium-mediated tobacco and maize transformation

The *LcCHI2* gene was amplified with the primers 5′-ccggaattcATGGCGAGGT TTGCTGCCCTCG-3′ and 5′-cgcggatccCTAGCTAGCGAAGTTTCGCTGGGTG-3′. The PCR products were digested with *Eco*R I / *Bam*H I and cloned into the pUC119-35s-NOS vector to generate pUC119-35s::*LcCHI2*-NOS. The fragment of 35s::*LcCHI2*-NOS was collected by *Hin*d III digestion and cloned into pBI121vector. After sequencing the positive clones, the final construct pBI121-35s∷*LcCHI2*-NOS were transformed into *Agrobacterium* strain EHA105. Transgenic tobacco plants were produced by using *Agrobacterium*-mediated transformation as described by the previously reported method (Hoekema *et al.*, 1983). Transgenic plants were selected on MS medium containing 50 mg/ml kanamycin and 500 mg/ml carbenicillin.

For maize transformation, the sequence of *LcCHI2* from pBI121-35s::*LcCHI2*-NOS *was* digested with *Spe*I / *Bam*H I and cloned into the pTF101-ubi-NOS vector to generate pTF101-ubi∷*LcCHI2*-NOS. Immature zygotic embryos of HiII (B73×A188) were infected with *Agrobacterium* strain EHA105 harboring the binary vector pTF101-ubi∷*LcCHI2*-NOS. Transgenic plants were selected and produced by the previously reported method (Frame *et al.*, 2002).

### Chitinase activity assay

Chitinase activity was measured in 10 mM potassium phosphate buffer (pH 7.0) containing 0.1 mM 4-methylumbelliferyl-*β*-D-*N*, *N’*-diacetylchitobiose (Sigma, Germany) as a substrate by the previously reported method (O’Brien and Colwell, 1987). One unit of chitinase activity was defined as the amount of enzyme required to produce 1 μmol of 4-methylumbelliferone per minute.

### Measurement of Malondialdehyde (MDA) contents and ion leakage ratio

Fresh leaves (0.3 g) were ground properly in 5 ml of 5 % trichloroacetic acid solution and centrifuged for 10 min at 3000 rpm. 2 ml of the supernatant was reacted with 2 ml 0.67% thiobarbituric acid and then it was heated for 30 min, at 100◻ in a water bath and then immediately cooled on ice. After centrifugation for 10 min, at 3000 rpm, the absorbance of the supernatant was read at 450, 532 and 600 nm. The contents of MDA were calculated using the formula: MDA concentration (μmol/L) = 6.45 × (A532-A600) − 0.56 × A450.

Ion leakage ratio was measured as relative electrical conductivity parameter. 0.1 g leaves were sampled from different plants, rinsed briefly with deionized water and immediately placed into a tube with 10 mL of deionized water. Conductivity (I1) was measured using an electroconductivity meter (model 1054, VWR Scientific, Phoenix) after the tubes were placed at 22◻ overnight. Then, the samples were heated at 100◻ for 30 min and conductivity (I2) was measured again. Relative electrical conductivity was expressed as (I1/I2) × 100%.

### Sodium content

Seedings of wild type (WT) and transgenic tobacco plants were cultured in nutrient solution for one month and then treated with 0 or 200 mM NaCl. After 6 days of treatment, the shoots and roots were oven dried, digested with H_2_SO_4_-H_2_O_2_.The digest was dissolved in deionised water and sodium content was estimated by flame spectrophotometer (M410, Waters Corporation, America).

### Salt stress treatments

The plant seedlings of Chinese wildrye (*Leymus Chinensis*) were grown in vermiculite with a light/dark cycle of 16 h/8 h at 25◻. They were irrigated with one-half-diluted Murashige and Skoog (MS) medium every 3 days. 4-week-old seedlings of Chinese wildrye were treated with 400 mM NaCl and 100 mM Na_2_CO_3_, respectively. Leaves of treated plants were harvested up to 0, 6, 12, and 24 h, respectively, by rapid freezing in liquid nitrogen and kept at -80◻ for further analysis.

To determine abiotic stresses tolerance of the transgenic tobacco (*Nicotiana tabacum*) on disc, seeds of wild-type and transgenic tobacco were sowed on MS medium containing 200 mM NaCl, 30 mM Na_2_CO_3_ or 500 mM sorbitol, respectively. For soil experiment, tobaccoseeds germinated in soil cultured with a light/dark cycle of 16 h/8 h at 25°C. Thirty plants for each line were used for salt and drought stress treatment, respectively. For salt stress, nearly 3-week-old plants were irrigated for 25 days with either tap water or tap water containing 400 mM NaCl or 100 mM Na_2_CO_3_.

To determine abiotic stresses tolerance of the transgenic maize, T2 generation transgenic event (CHI-Ox2, CHI-Ox5 and CHI-Ox7) and WT plants were grown in pots (10×10 cm) containing peat: vermiculite (5:1, v/v) medium in a greenhouse with a light/dark cycle of 16 h/8 h at 30°C. PCR analysis was used to screen for positive transformants, and 2-week-old seedlings were water with either tap water or tap water containing 200 mM NaCl or 50 mM Na_2_CO_3_ for 6 days. Photograph was recorded after each treatment by a camera.

### Pathogen response assays of transgenic tobacco and Maize

For bacterial infection analysis, *Pseudomonas tabaci (Wolf et Foster) Stevens* was grown in NBY medium and bacterial cells were collected, washed and resuspended in 10 mM MgSO_4_. The density of bacterial cells was determined by measuring absorbance at OD_600_. Bacterial cells in suspension were infiltrated into fully expanded 6-week-old tobacco leaves using a 1 ml plastic syringe without a needle. After 5 days, the average lesion area for each independent transgenic line was calculated and compared with that of wild-type tobaccos.

For fungal resistance analysis, *Alternaria alternata (Fries) Keissler* was cultured on potato dextrose agar medium at 28◻. When the mycelia reached the edge of the plate, 0.5 cm diameter agar discs were excised from the edge of growing colonies using a cork borer and inverted onto the detached leaves from wild-type and transgenic tobaccos. All leaves were placed on wet filter paper in Petri dishes and incubated at 28◻ to permit normal disease development under high humidity. After 7 days, the average lesion area for each independent transgenic line was calculated and compared with that of wild-type tobaccos.

Maize pathogen *Exserohilum turcicum* and *curvularia lunata* were provided by Iinstitute of plant protection, Jilin academy of agricultural sciences. *Exserohilum turcicum* and *curvularia lunata* were cultured on potato dextrose agar medium at 28◻. When the mycelia reached the edge of the plate, 0.6 cm diameter agar discs were excised from the edge of growing colonies using a cork borer and inverted onto the detached leaves from transgenic and null control maize plants. All leaves were placed on wet filter paper in Petri dishes and incubated at 28◻ to permit normal disease development under high humidity. After 3 days, the average lesion area for each independent transgenic line was calculated.

### Nucleotide and protein sequences accession numbers

The nucleotide and protein sequences reported in this paper have been deposited in the Genbank nucleotide database and protein database under the accession number GQ397277 and ACV20870.

## Results

### Cloning and characterization of *LcCHI2* from Chinese wildrye

According to the EST sequence information from literature (Jin *et al.*, 2006), we cloned a full-length cDNA of the target chitinase gene by 3‘RACE technology from Chinese wildrye, and was designated as *LcCHI2* (GenBank accession number GQ397277). Sequence analysis revealed that *LcCHI2* contains a 771 bp open reading frame encoding a polypeptide of 256 amino acids. Pfam scan showed that the deduced protein is a class II chitinase belonging to the GH19 family (PF00182) (https://www.ebi.ac.uk/Tools/pfa/pfamscan/). To obtain more information of the LcCHI2 protein, a phylogenetic tree was constructed including LcCHI2 orthologes and all the class II chitinases of Arabidopsis. LcCHI2 belonged to a cluster including three pathogenesis-related protein 3 (HvPR3, TaPR3 and AtPR3) and two chitinases (ScCHI46, CaCHI2) (Figure S1). LcCHI2 showed more than 95% homology to HvPR3, TaPR3 and ScCHI46 and 71% homology to CaCHI2 (Figure S2A). Multiple protein alignment revealed two conserved chitinase family 19 signature domains in all the five orthologes (Figure S2B). However, an *N*-terminal secretive signal peptide only found in LcCHI2 and its monocot orthologes (Figure S2B).

### Sodium stress induced the expression of *LcCHI2* and chitinase activity in Chinese wildrye

As the EST of *LcCHI2* was isolated from alkaline-sodic stressed tissues, we measured the expression of *LcCHI2* in Chinese wildrye under different abiotic conditions. A nearly 8-fold induction of *LcCHI2* transcripts was observed under 400 mM NaCl after 24 hours treatment comparing with the control plant (Figure 1A). Similarly, the expression of *LcCHI2* increased more than 2-fold under 100 mM Na_2_CO_3_ after 12 and 24 hours treatment compared with the control plant (Figure 1A). Subsequently, the chitinase activities were determined under NaCl and Na_2_CO_3_ conditions after 24 hours treatment. The results showed that chitinase activities increased by 4.3-fold and 2.6-fold after 24 hours exposure to 400 mM NaCl and 100 mM Na_2_CO_3_, respectively (Figure 1B). In contrast, the expression of *LcCHI2* and chitinase activity were not changed in Chinese wildrye under 20% PEG condition in 24 hours (data not show).

**Figure 1.**
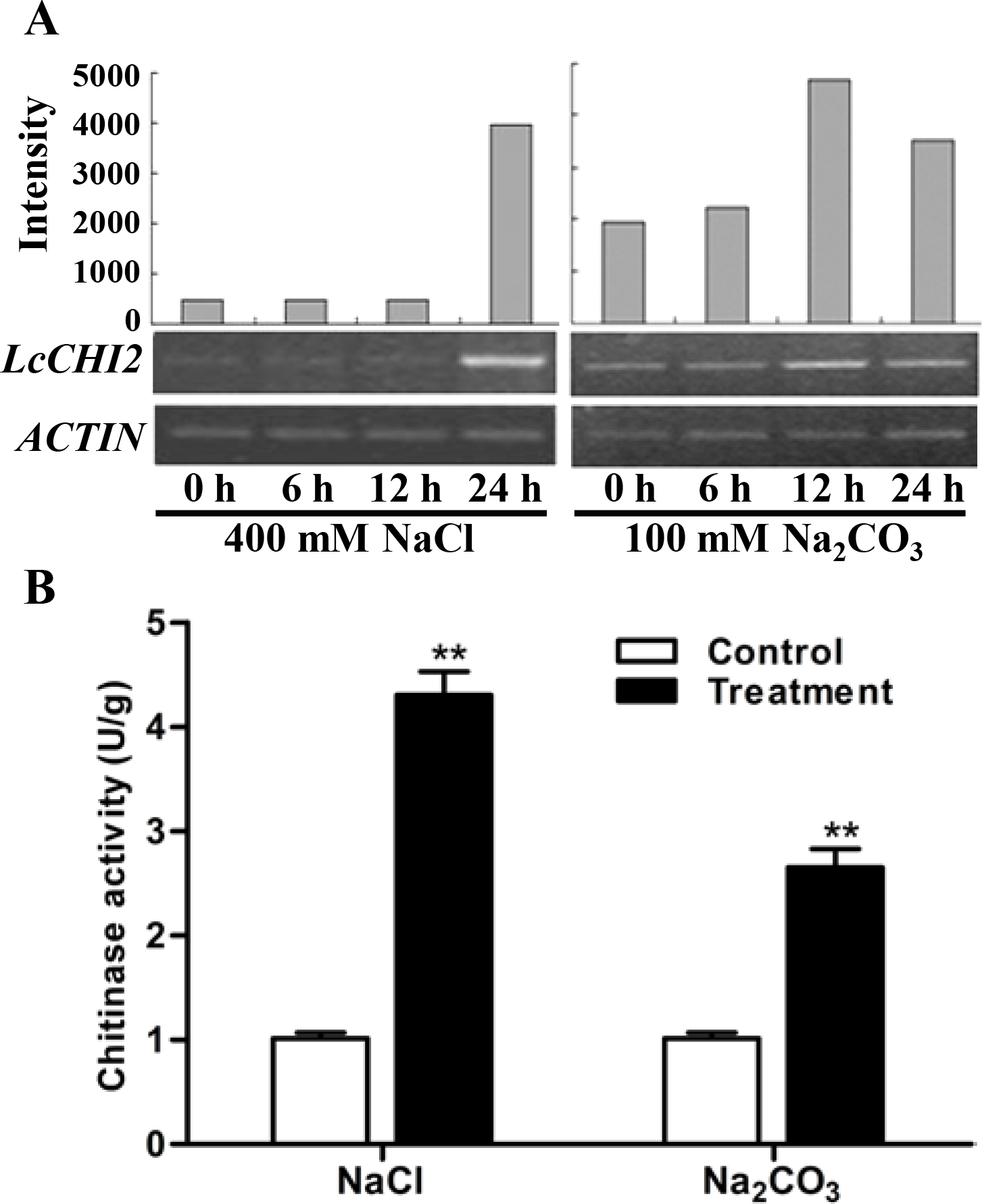
Regulation of *LcCHI2* and chitinase activity under salt stress conditions in Chinese wildrye. (A) Expression analysis of the *LcCHI2* after treatment with 400 mM NaCl and 100 mM Na_2_CO_3_ for the indicated times. Total RNA were extracted from 4-week-old seedlings of Chinese wildrye following treatments as indicated and reverse transcribed. The cDNA were used as templates for RT-PCR and the *ACTIN* gene was amplified as an internal control. The PCR products were examined by electrophoresis in 1% (w/v) agarose gel. (B) Chitinase activity assay under salt stress conditions in Chinese wildrye. Enzyme activity assays were carried out with the leaves of Chinese wildrye after treatment with 400 mM NaCl and 100 mM Na_2_CO_3_ for 24 hours. Values are the mean ± SE obtained from three biological replicates. ** one-way ANOVA; P < 0.01.

### Overexpression of *LcCHI2* in tobacco enhanced plant tolerance to salt stress

To further investigate the function of *LcCHI2*, transgenic tobacco plants that constitutively express the *LcCHI2* gene under the control of the CaMV 35S promoter were developed by agrobacterium-mediated transformation. Three independent positive transgenic lines were selected by kanamycin resistance and used for further analysis. The expression of *LcCHI2* could be detected in all the three transgenic lines other than wild-type (WT) plants (Figure 2A). The results showed that chitinase activity increased by 1.5 - 1.7 folds over WT plants in the transgenic lines (Figure 2B). Comparing with the WT plants, the *LcCHI2*-overexpression plants exhibited no obvious phenotypic difference from the WTs under normal growth conditions (Figure S3).

**Figure 2.**
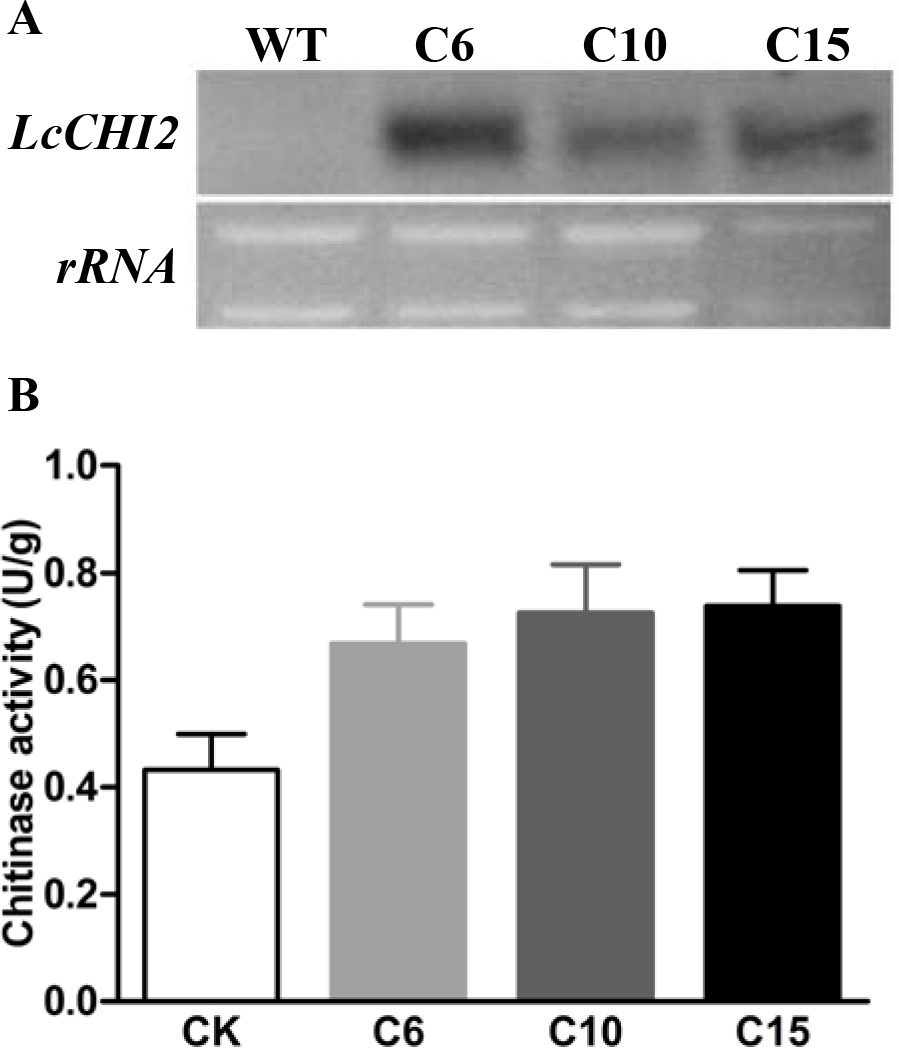
(A) Northern blot analysis of *LcCHI2* transcripts in wild-type (WT) and transgenic tobacco plants (C6, C10, C15). A DIG-labeled LcCHI2 probe was used for hybridization. (B) Chitinase activity of WT and transgenic tobacco plants. Values are the mean ± SE obtained from three biological replicates.

As the expression of *LcCHI2* is induced by salt stress, phenotypes of the WT and *LcCHI2*-overexpression lines were analyzed on MS medium containing 200 mM NaCl, 30 mM Na_2_CO_3_ and 500 mM sorbitol, respectively. No difference was observed between WT and *LcCHI2*-overexpression plants growth on normal MS medium. However, the *LcCHI2*-overexpression plants survived well under salt stress (NaCl and Na_2_CO_3_) in comparison with WT plants, especially the C10 and C15 lines (Figure 3A). In contrast, the *LcCHI2*-overexpression plants only exhibited a minor effect for osmotic stress (Figure 3A). To further evaluate the abiotic resistance of *LcCHI2*-overexpression plants, soil growth plants were growth under salt and drought stress conditions. There were no observed differences in phenotypes between *LcCHI2*-overexpression and WT plants, when they were growth under normal and drought stress conditions (Figure S4). However, 6 days after watering 400 mM NaCl or 100 mM Na_2_CO_3_, the wild-type seedlings showed wilting phenotypes and ultimately died, whereas the transgenic tobaccos continued to grow (Figure S4).

**Figure 3.**
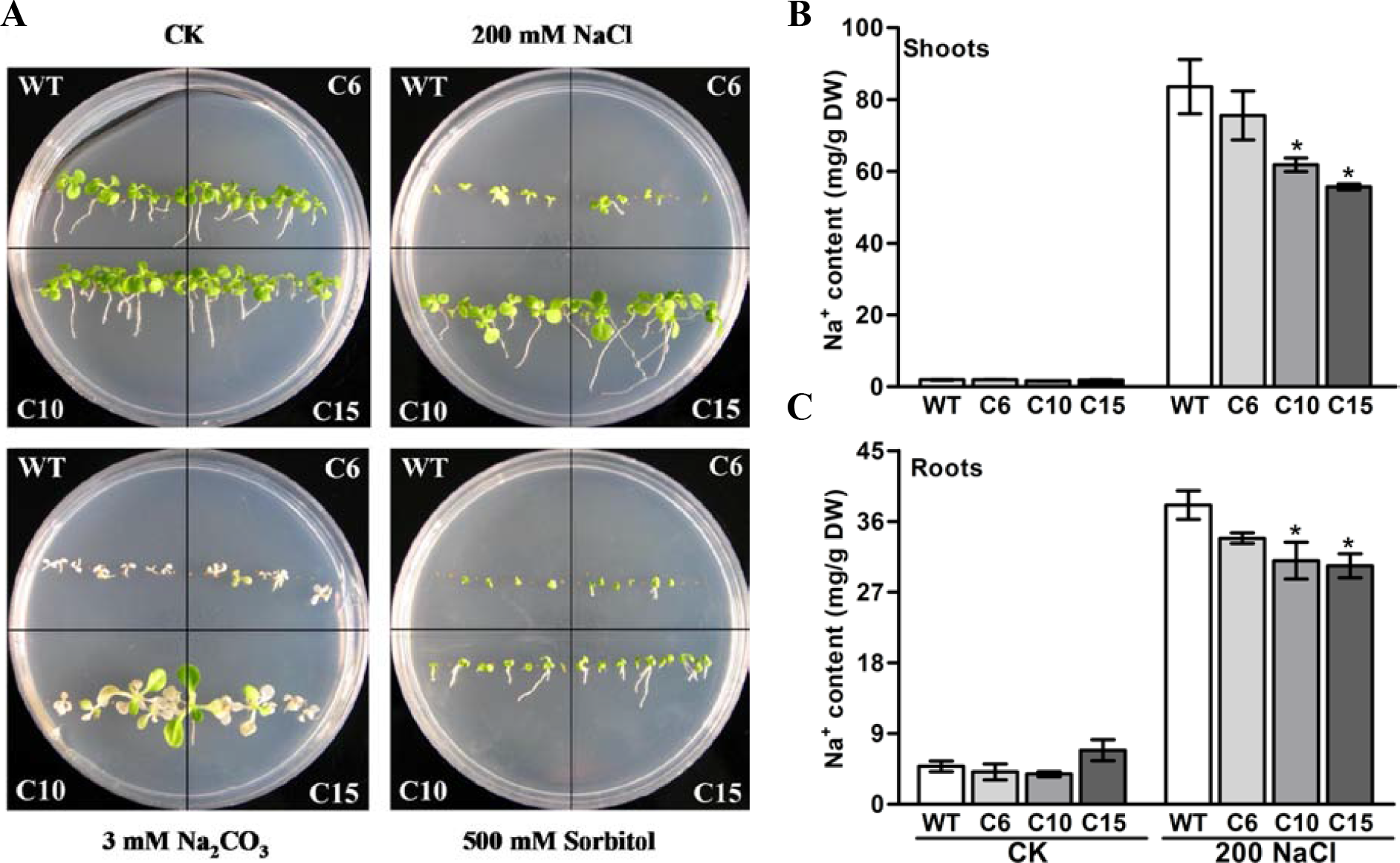
Abiotic stress tolerance assays of wild-type and *LcCHI2*-overexpressed plants in tobacco. (A) The WT and *LcCHI2* overexpression (C6, C10 and C15) plants were grown on MS agar plates containing 200 mM NaCl, 30 mM Na_2_CO_3_ and 500 mM sorbitol for 30 days. (B) and (C) Na^+^ content of wild-type and *LcCHI2* overexpression plants in shoots and roots. Values are the mean ± SE obtained from three biological replicates.

Because the overexpression of *LcCHI2* increased salt stress tolerance other than osmotic or drought stresses, sodium contents were determined in the roots and shoots of WT and transgenic plants under normal and NaCl treatment conditions. As expected, NaCl treatment significantly increased Na^+^ content in the shoots and roots of WT and transgenic plants (Figure 3B, 3C). The overexpression of *LcCHI2* decreased the Na^+^ content from 10% to 33% in the shoots and from 11% to 22% in the roots of transgenic lines compared with the corresponding WTs under NaCl treatment condition (Figure 3B, 3C). Meanwhile, the Na^+^ content showed a significantly decrease only in transgenic lines C10 and C15 other than line C6, which is coincident with the strong resistance to sodium stress of the transgenic lines C10 and C15 (Figure 3A).

Malondialdehyde (MDA) content and relative electrical conductivity are widely used as indicators for lipid peroxidation and the degree of plant cell injury under stress treatment, respectively. There are no obvious differences in MDA content and relative electrical conductivity between WT and *LcCHI2* overexpression plants under normal growth condition. Although 24 hours salt treatment significantly increased the MDA content and relative electrical conductivity in both WT and *LcCHI2* overexpression plants, the transgenic lines showed a lower MDA content and relative electrical conductivity compared with WT under salt stress condition (Figure S5).

### Overexpression of *LcCHI2* conferred pathogen tolerance in tobacco

As a homologue of PR3 proteins, LcCHI2 is presumed to be involve in pathogen resistance too. Thus, the *LcCHI2*-overexpression plants were inoculated with bacterial and fungal pathogens, respectively. Leaves of WT and *LcCHI2*-overexpression tobaccos were inject-inoculated with spore suspensions of the bacterial pathogen *Pseudomonas tabaci (Wolf et Foster) Stevens* and symptom development was subsequently monitored for 5 days. The examined disease symptoms included chlorosis and necrosis expansion surrounding the primary infection sites. As shown in Fig. 4, WT tobaccos were more sensitive to bacterial infection than *LcCHI2*-overexpression tobaccos in detached leaves, in which pathogen infection resulted in significantly reduced disease symptoms. The leaf lesion size of WT tobaccos was 5- to 10-fold larger than that of *LcCHI2*-overexpression tobaccos (Figure. 4A, 4B). Resistance to the fungal pathogen *Alternaria alternata (Fries) Keissler* was also identified in *LcCHI2*-overexpression tobacco plants by a detached leaf inoculation test after 7 days inoculation. WT tobaccos showed typical necrosis symptoms surrounded by chlorotic halos and extensive pathogen sporulation, while transgenic lines had significantly smaller lesion sizes than wild-type tobacco (Figure. 4C, 4D). The enhanced resistance to *Pseudomonas tabaci and Alternaria alternata* in transgenic tobacco demonstrated that disease resistance conferred by *LcCHI2* overexpression is effective against both bacterial and fungal pathogen.

**Figure 4.**
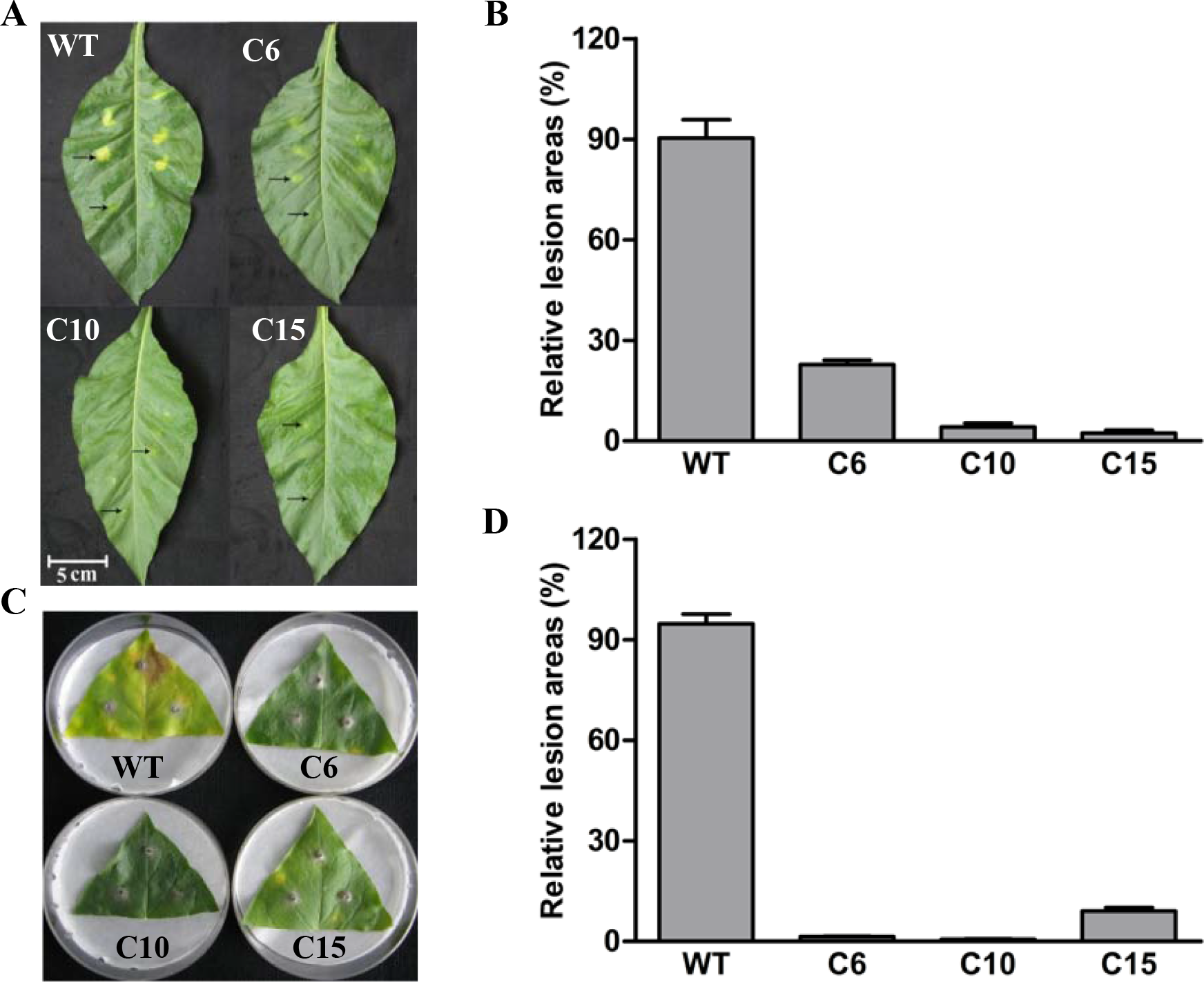
Pathogenesis analysis of *LcCHI2*-transgenic tobacco. (A) and (B) Resistance of transgenic tobacco to the bacterial pathogen *Pseudomonas tabaci* (*Wolf and Foster*) *Stevens*. Fully expanded leaves of tobaccos were syringe-infiltrated with 0.1 ml solution of*P. tabaci*. Arrows in the upper line represent the sites inoculated with bacteria and arrows in the lower line represent the sites with mock-inoculated. The average lesion area of each independent transgenic line was calculated and their relative lesion areas are shown in columns after comparison with the average lesion area on wild-type tobacco. The photograph was taken 5 days after inoculation. (C) and (D) Responses of transgenic tobacco to the fungal pathogen *Alternaria alternata (Fries) Keissler*. Detached leaves were challenged with mycelia of *A. alternata*. The average lesion area of each independent transgenic line was calculated and their relative lesion areas are shown in columns after comparison with the average lesion area on wild-type tobacco. The photograph was taken 7 days after inoculation. Values are the mean ± SE obtained from three biological replicates.

### Overexpression of *LcCHI2* in Maize enhanced tolerance to salt and pathogen stresses

To determine whether the overexpression of the *LcCHI2* gene could phenocopy the stress tolerances of transgenic tobacco in monocotyledons crops, we produced transgenic maize plants harboring the *pTF101.1-ubi∷LcCHI2* construct. DNAs were extracted from leaves of T_0_ generation transgenic plants and use to detect the integration of target sequence into genomes of transgenic plants by PCR. The results showed that six transgenic events harbored both the selection marker and *LcCHI2* genes (Figure S6). Reverse-transcriptional PCR (RT-PCR) and chitinase activity assay indicated that three representative transgenic lines accumulated abundance *LcCHI2* transcripts and enhanced chitinase activities compared to null plants with the absence of *LcCHI2* (Figure 5A, 5B). Then these transgenic plants were grown in normal and alkaline soils in the greenhouse (Figure S7). There are no significant different between the growth of transgenic (Ox) and their respective non-transgenic plants in normal soil (null). However, the transgenic plants were healthier than the corresponding null plants in alkaline soil. A significant increased plant height and SPAD value were observed in transgenic plants compared their null plants (Figure S7B, S7C). To further evaluate the effects of divergent salt stress on the transgenic maize, two week-old plants were watered with 200 mM NaCl and 50 mM Na_2_CO_3_, respectively. As expected, no significant differences were observed between the growths of transgenic (Ox) and their respective null plants after six-day treatment with distilled water (Figure 6A, 6B). In contrast, the overexpression plants showed resistance to NaCl or Na_2_CO_3_ treatments other than the null plants (Figure 6C, 6D). Although NaCl or Na_2_CO_3_ treatments increased the relative electrical conductivity and MDA content in both overexpression and null plants, the magnitudes of the overexpression plants were significantly lower than null plants (Figure 6E, 6F).

**Figure 5.**
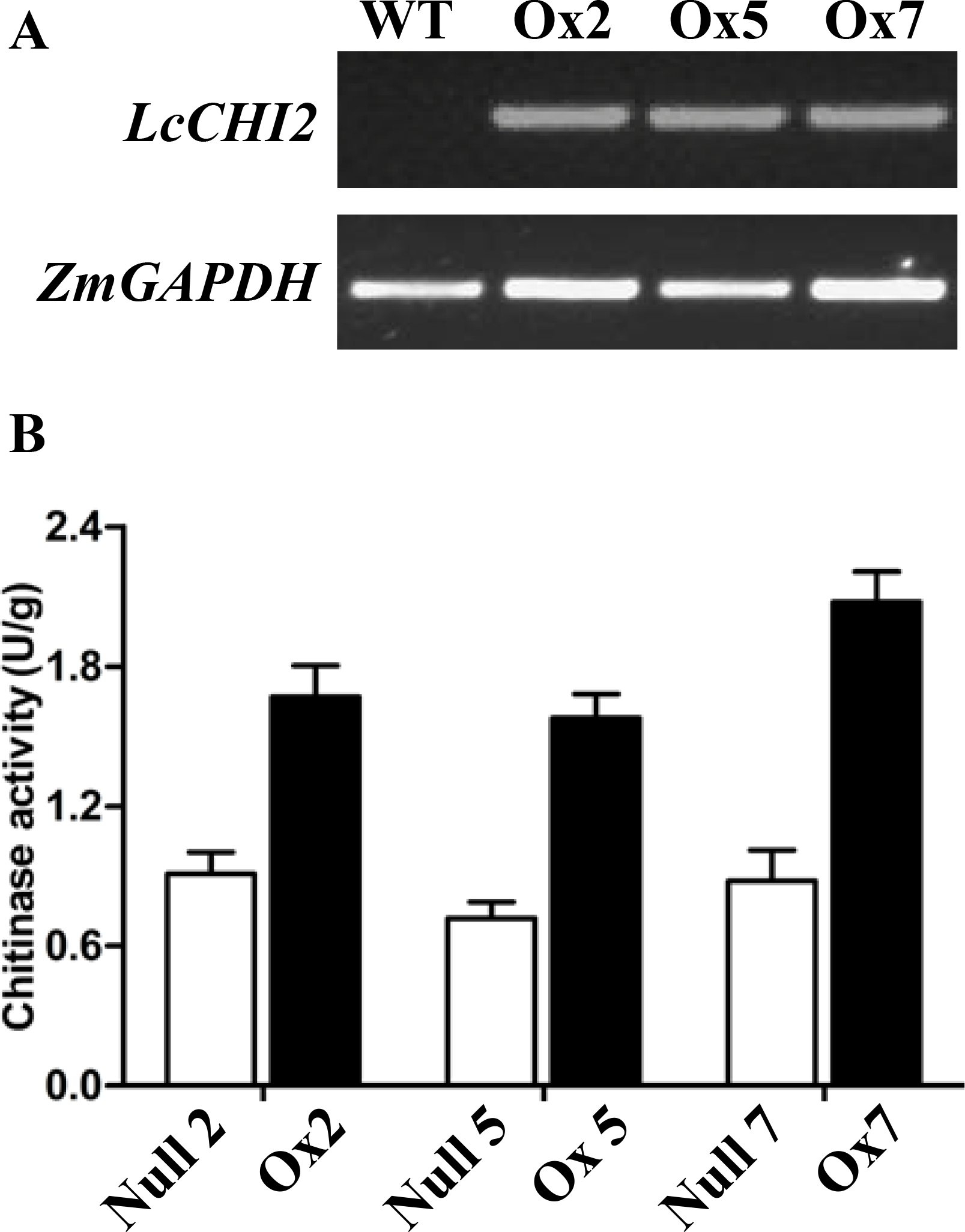
(A) RT-PCR analysis of *LcCHI2* transcript in wild-type (WT) and transgenic Maize plants (Ox2, Ox5, Ox7). Total RNA were extracted from 2-week-old seedlings of positive and null transgenic plants. The cDNA were used as templates for RT-PCR and the actin gene was amplified as an internal control. The PCR products were examined by electrophoresis in 1% (w/v) agarose gel. (B) Chitinase activity of null and transgenic maize plants. Values are the mean ± SE obtained from three biological replicates.

**Figure 6.**
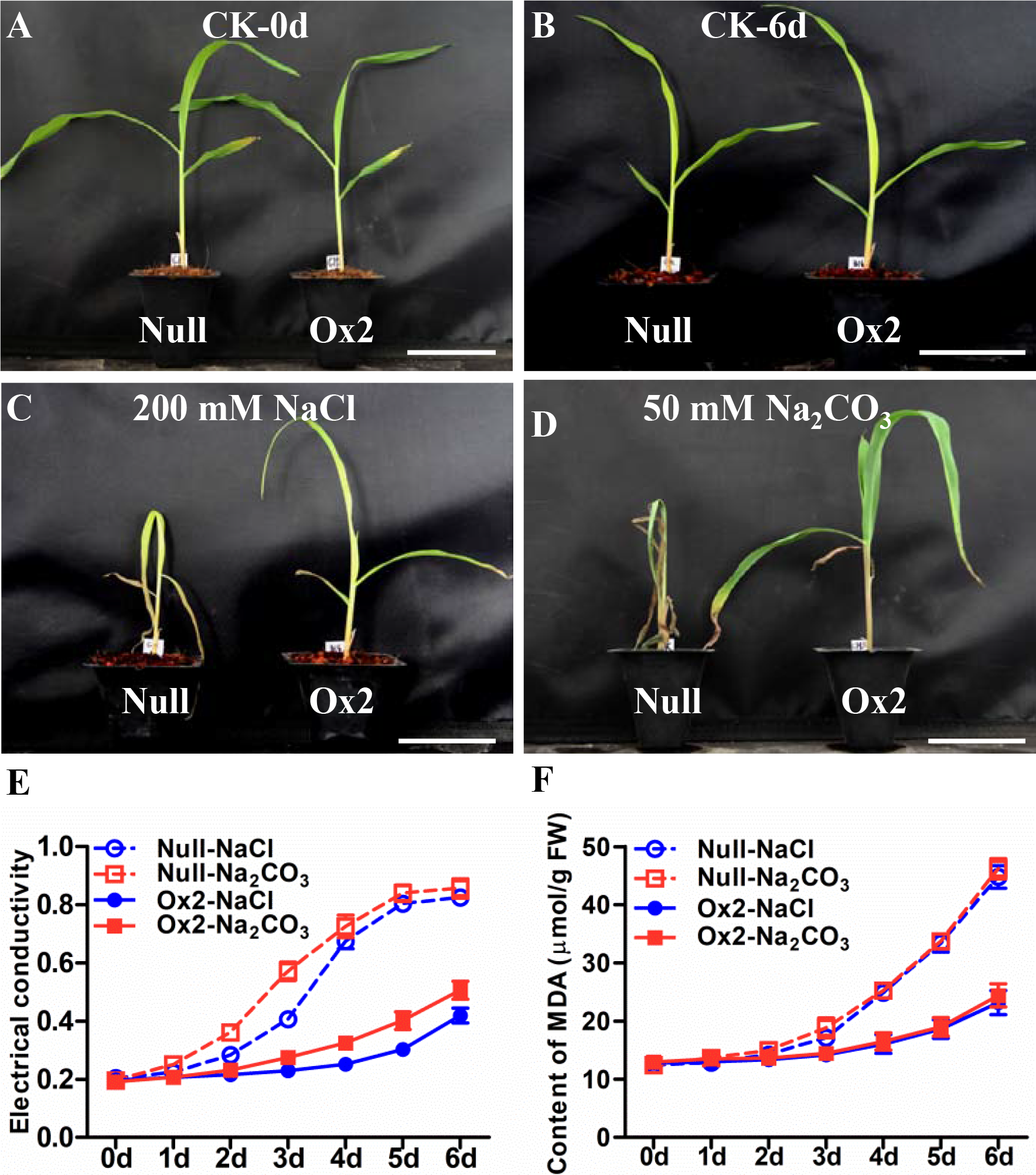
Salt stress tolerance assays of null and *LcCHI2* overexpression plants in maize. (A), (B), (C) and (D) Soil growth null and *LcCHI2* overexpression plants were watered with tap water, 200 mM NaCl and 50 mM Na_2_CO_3_ for 6 days. (E) and (F) MDA contents and relative electrical conductivity of null and *LcCHI2* transgenic maize were measured every day. Values are the mean ± SE obtained from three biological replicates. * one-way ANOVA; P < 0.05.

Biotic resistances of the transgenic plants were also measured by infection the maize with two major fungal pathogens *Exserohilum turcicum* and *curvularia lunata*, respectively. Typical necrosis symptoms were observed in the null plants while transgenic lines had significantly smaller lesion area after infection by *Exserohilum turcicum* or *curvularia lunata* (Figure. 7A, 7B, 7C). These results indicated that the overexpression of *LcCHI2* in maize significantly inhibited the growth of fungal pathogens.

**Figure 7.**
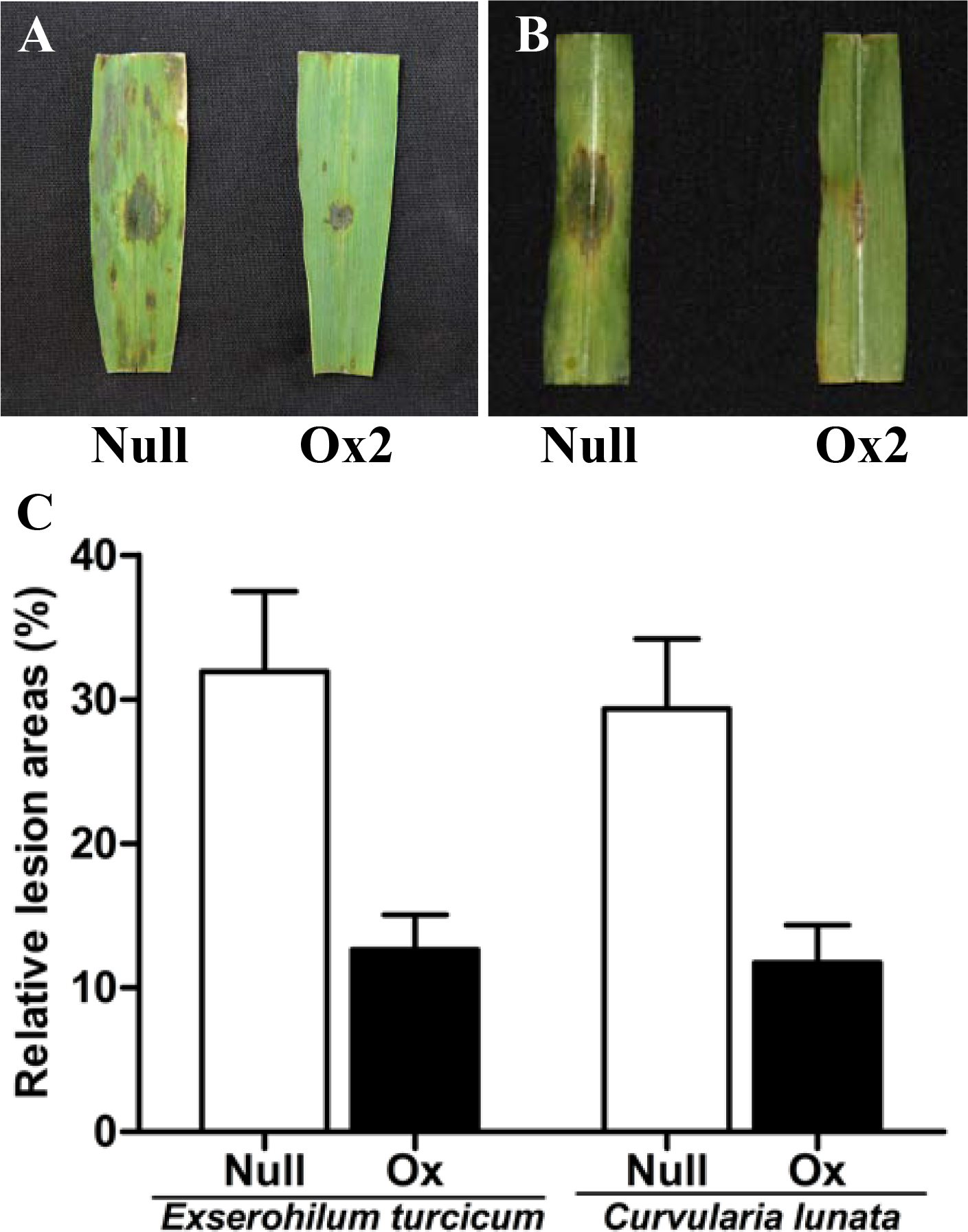
Pathogenesis analysis of null and *LcCHI2*-overexpressed maize. (A) Resistance of transgenic maize to fungal pathogen *Exserohilum turcicum*. (B) Responses of transgenic maize to the fungal pathogen *curvularia lunata*. (C) The average lesion area of each independent transgenic line was calculated and their relative lesion areas are shown in columns after comparison with the average lesion area on wild-type tobacco. Values are the mean ± SE obtained from three biological replicates.

To evaluate the effects of *LcCHI2* overexpression in agronomic traits, the gene in the transgenic events with HiII background (CHI-Ox2, CHI-Ox5 and CHI-Ox7) were backcross to local commercial inbred line X923-1. The genetically segregated positive and negative BC5F2 plants were crossed respectively with Y822 to produce hybrid seeds. Agronomic traits were measured from the hybrid lines in two replicated experiments. The results (Table S1) showed that all transgenic events exhibited statistically no negative impact such as cob weight, cob diameter, ear kernel weight and ear diameter, indicating that overexpression of *LcCHI2* has no penalty on the growth and yield in the field conditions comparing with the controls.

## Discussion

In this study, the full-length cDNA of a chitinase-like gene was cloned and named as *LcCHI2*, which was first identified from an EST library in *Leymus Chinensis* under Na_2_CO_3_ stress (Jin *et al.*, 2006). The cDNA-deduced protein sequence analysis indicated that the LcCHI2 contains two conserved domains including the signature 1 (PS00773) and signature 2 (PS00774) of glycosyl hydrolase 19 family (Figure S2B). Protein cluster analysis indicates that LcCHI2 belongs to class II chitinase (Figure S1). Several members of the class II chitinase proteins, such as AtPR3 and pcht28, were reported to have the typical chitinase activity and overexpression of those chitinase genes could increase chitinase activity in transgenic plants (Verburg and Huynh, 1991; Tabaeizadeh *et al.*, 1999; Chen *et al.*, 1994). In this study, a significantly increased chitinase activity was positively correlated with the up-regulation of *LcCHI2* in *Leymus Chinensis* under both NaCl and Na_2_CO_3_ stress condition. Moreover, the transgenic tobacco and maize plants that overexpressed *LcCHI2* also showed higher chitinase activity than control plants. Interestingly, protein alignment indicated that LcCHI2 and its homologues contain an N terminal secretary signal peptide (Figure S2B). These results supported that LcCHI2 is a secreted active chitinase in *Leymus Chinensis*.

Different types of chitinases were induced by differential pathogen attack and have been characterized as pathogenesis-related (PR) proteins (Kesari *et al.*, 2015; van Loon *et al.*, 2006). Phylogenic analysis showed that LcCHI2 is a homologue of several reported PR3 proteins, including TaPR3, HvPR3, CaCHI2, pcht28 and AtPR3 (Figure S2) (Verburg and Huynh, 1991; Hong and Hwang, 2002; Tabaeizadeh *et al.*, 1999; Shin *et al.*, 2008; Scheler *et al.*, 2016). These PR3 proteins were proposed to be involved in plant defense responses. Plant chitinases were thought to degrade the major structural polysaccharide of fungal cell walls in the intercellular space to limit fungal growth (Brogue *et al.*, 1991). The purified AtPR3 protein showed antifungal chitinase activity and inhibited the growth of *Trichoderma reesei in vitro* (Verburg and Huynh, 1991). The overexpression of chitinase resulted in enhanced resistance in tomato, rice, Italian ryegrass and grapevine to *Verticillium dahlia*., Magnaporthe *grisea*, *Puccinia coronata* and *Uncinula necator*, respectively (Yamamoto *et al.*, 2000; Tabaeizadeh *et al.*, 1999; Nishizawa *et al.*, 1999; Takahashi *et al.*, 2005). Moreover, expression of a barley *PR3* chitinase gene in transgenic wheat resulted in enhanced resistance to infection by *Erysiphe graminis*, *Blumeria graminis*, *Pucinia recondita* and *Fusarium graminearum* (Shin *et al.*, 2008; Oldach *et al.*, 2001). To elucidate the defense response of *LcCHI2*, we investigated the biological functions of the *LcCHI2* in transgenic tobacco and maize. The overexpression of *LcCHI2* significantly enhanced resistance to multiple pathogens in tobacco and maize. The transgenic tobacco plants also showed resistance to a bacterial pathogen *Pseudomonas tabaci Stevens*, which do not have chitin or related structure in their cell wall. A similar phenomenon was observed in the transgenic Arabidopsis when overexpressing *CaCHI2* gene (Hong and Hwang, 2006). In fact, chitinase expression in plant tissues is strongly induced by infection with viruses and bacteria in addition to fungal pathogen (van Loon *et al.*, 2006). Therefore, the PR3 type protein may inhibit the pathogen invasion by an unknown mechanism other than directly degrading the chitin oligomers.

In addition to biotic stress, a number of cluster II chitinases were reported to be involved in developmental and various abiotic responses. The ScCHT46 showed 92% protein identify with LcCHI2 and encode a chitinase-antifreeze protein in winter rye (Yeh *et al.*, 2000). The expression of *ScCHT46* and its homologs in winter wheat, *HvPR3*, are responsive to cold and drought (Yeh *et al.*, 2000). Moreover, it is reported that NaCl, drought and Mannitol induces the expression of several class II chitinases, including *CaCHI2*, *pcht28* and *AtPR3* (Seo *et al.*, 2008; Hong and Hwang, 2002, 2006; Chen *et al.*, 1994). The overexpression of *CaCHI2* increased the osmotic stress in Arabidopsis (Hong and Hwang, 2006). In contrast, a mutation of *AtPR3* affected the seeds geminating under high salt (Seo *et al.*, 2008). Similarly, our results showed that the expression of *LcCHI2* was up-regulated by Na^+^ stress conditions including treatment with NaCl and Na_2_CO_3_. Moreover, the activity of chitinase was increased under NaCl and Na_2_CO_3_ stresses in Chinese wildrye. Taking together, these results suggested that *LcCHI2* play an important role in abiotic stress tolerance similar to its homologues in other plants.

Interestingly, the overexpression of *LcCHI2* showed a significant tolerance to NaCl and Na_2_CO_3_ treatments in both tobacco and maize but not to the osmotic or drought stresses. These results implicated that the overexpression of *LcCHI2* might assist to reduce Na^+^ ionic toxicity other than osmotic stress or improving water-use-efficiency stress. Coincidence with this assumption, the Na^+^ contents were decreased in the transgenic tobacco under NaCl treatment. Recently, it is proposed that the amount of the carboxyl groups on de-methesterified pectin could bind cations such as Na^+^ and sequester more Na^+^ in the cell wall, which may contribute to salt stress resistance in plants (de Lima *et al.*, 2014; Byrt *et al.*, 2018; An *et al.*, 2014). However, we did not detect any significant difference in the pectin content between transgenic and WT plants (data not show). Instead, increased cellulose and decreased hemicelluloses were observed in the overexpression lines compared with WT plants under salt treatment condition (Fig S8). This phenomenon is reminiscent of the Na^+^ sensitive mutant *Atctl1*, which affects the contents of cellulose and extractability of hemicelluloses (Sanchez-Rodriguez *et al.*, 2012; Hermans *et al.*, 2011). It is reported that mutation of *AtCTL1* increased the Na^+^ influx rather than defects in Na^+^ efflux activity (Kwon *et al.*, 2007). Therefore, the overexpression of the *LcCHI2*, a homologue of AtCTL1 (Fig S2), may have an opposite effect and inhibit the Na^+^ influx, which could explain the deceased Na^+^ content in *LcCHI2* transgenic plants.

Although cell wall metabolism is rationally correlated with salt stress, the exact mechanism is still largely unknown (Le Gall *et al.*, 2015). Mutation of *AtCES8* (cellulose synthase 8) or *AtCSLD5* (cellulose synthase-like 5) resulted tolerance or hypersensitive to salt stress, respectively (Zhu *et al.*, 2010; Chen *et al.*, 2005). Moreover, mutation the chitinase-like gene *AtCTL1* reduced the cellulose content and led to salt-sensitive of Arabidopsis (Sanchez-Rodriguez *et al.*, 2012; Kwon *et al.*, 2007). Compared with dicot, there has been no report on the modification of cell wall compound contributing to salt tolerance in grass. In this study, the overexpression of *LiCHI2* in both tobacco and maize enhanced the salt and pathogen stress tolerances, suggesting similar roles of the chitinase in dicot and grass. Although the overall architectures of plant cell walls are similar in that they both consist of a network of cellulose fibers surrounded by a matrix of non-cellulosic polysaccharides, the types and abundance of non-cellulosic polysaccharides are significant different between grass and dicot cell walls (Vogel, 2008). It seems that cellulose represents the common component in plant cell walls and plays a pivotal role in regulating the salt stress. The changed cellulose content may regulate cross-linking and rigidity of the cell wall, which acts as a barrier for salt entrance or pathogen invasion.

Although a large number of alkali-responsive genes were reported from different plants, a few have been cloned and characterized (Zhao *et al.*, 2016). Overexpression of the alkali-stress-inducable RMtATP6 and NADP-ME_2_ in rice and Arabidopsis enhanced the tolerance against osmotic, NaCl and Na_2_CO_3_ stresses (Liu *et al.*, 2007; Zhang *et al.*, 2006). Transgenic rice with *PtNHA1* and *PutNHX* genes, which were cloned from an alkaline soil plant, increased tolerance of shoots to NaCl and roots to NaHCO_3_, respectively (Kobayashi *et al.*, 2012). It is noted that all these studies were conducted in nutrient solution or agar plate in laboratory. In contrast, we presented the performance of the transgenic maize with natural alkaline soil or treated soil (Figure S7). Moreover, the transgenic maize showed no penalty on growth and yield in field condition (Table S1). Therefore, we proposed that overexpression of *LcCHI2* represent a potential strategy for engineering biotic and alkaline resistance plants.

## Supplementary data

Fig. S1 Phylogenetic tree constructed with class II chitinase sequences retrieved by BLAST in the NCBI database.

Fig. S2 (A) Protein similarity of LcCHI2 with its homologues. (B) Alignment of deduced amino acid sequences of LcCHI2 with its homologues from other plant species.

Fig. S3 Morphology of the *LcCHI2*-overexpressing transgenic tobacco plants under normal growth condition.

Fig. S4 Drought and salt stress tolerance assays of wild-type and LcCHI2 overexpression plants grow in soil

Fig. S5 MDA contents and relative electrical conductivity of wild-type and *LcCHI2*-transgenic tobaccos under 200 mM NaCl treatment.

Fig. S6 PCR detection of selection marker (a) and *LcCHI2* genes (b) in transgenic maize plants.

Fig. S7 Growth assessment of transgenic maize in normal and alkaline soils.

Fig. S8 Cellulose and hemicelluloses contents in wild-type and *LcCHI2*-transgenic tobaccos under 200 mM NaCl treatment.

Table S1. Agronomic traits of transgenic maize grow in the field condition.

## Acknowledgments

We thank Dr Qiyun Li for the help in setting up the experiments of pathogen response assays of transgenic tobacco. We thank Dr Bo Xu for the correction of the manuscript. This research was supported by the National Science Foundation of China (31772378, 31771879), the Science and Technology Innovation Program of Jilin Province (CXGC2017JQ014) and Huazhong Agricultural University Scientific & Technological Self-innovation Foundation (2016RC005, 2662017PY012).

